# The ABC transporter Snu drives formation of the lipid-based inward and outward barrier in the skin of *Drosophila*

**DOI:** 10.1101/169466

**Authors:** Renata Zuber, Michaela Norum, Yiwen Wang, Kathrin Oehl, Davide Accardi, Slawomir Bartozsewski, Nicole Gehring, Jürgen Berger, Matthias Flötenmeyer, Bernard Moussian

## Abstract

Lipids in extracellular matrices (ECM) contribute to barrier function and stability of epithelial tissues such as the pulmonary alveoli and the skin. In insects, skin waterproofness depends on the outermost layer of the extracellular cuticle termed envelope that contains cuticulin, an unidentified water-repellent complex molecule composed of proteins, lipids and catecholamines. Based on live-imaging analyses of fruit fly larvae, we find that initially envelope units are assembled within vesicles harbouring the ABC transporter Snu and the extracellular protein Snsl. In a second step, the content of these vesicles is distributed to cuticular lipid-transporting nanotubes named pore canals and to the cuticle surface in dependence of Snu function. Consistently, the surface of *snu* and *snsl* mutant larvae is depleted from lipids and cuticulin. By consequence, these animals suffer uncontrolled water loss and penetration of xenobiotics. Our data allude to a two-step model of envelope i.e. barrier formation. The proposed mechanism in principle parallels the events occurring during differentiation of the lipid-based ECM by keratinocytes in the vertebrate skin suggesting establishment of analogous mechanisms of skin barrier formation in vertebrates and invertebrates.

## Introduction

Commonly, extracellular matrices (ECM) play essential roles in epithelial tissue function. Compared to proteins and sugars, lipids as constituents of ECMs have been studied only in few cases. In vertebrates, for instance, the outermost protective layer of the skin consists of a lipid-based ECM, the lamellar membrane, a network of lipids and proteins produced by keratinocytes (Koster, 2009; Nishifuji and Yoon, 2013). Mutations abrogating this matrix are associated with severe skin diseases in humans (Roberson and Bowcock, 2010; van Smeden et al., 2014). Another prominent example of a lipid-based ECM is the pulmonary surfactant that stabilises alveoli (Goerke, 1998). Failure to produce the surfactant provokes lethal lung diseases (Beers and Mulugeta, 2017). In insects, lipids are reported to cover the body surface and to impregnate the non-cellular cuticle, which is assembled at the apical side of the epidermis (Moussian, 2010). Surface lipids, mostly alkanes and alkenes, are free, and have been considered to play an important role in dehydration and penetration protection (Gibbs, 1998; Gibbs, 2002; Wang et al., 2016; Wang et al., 2017). Integral lipids are immobilised within the cuticle through association with proteins and catceholamines. These complex molecules that besides of being key components of the waterproof barrier, also contribute to cuticle colour and stiffness, were named cuticulin by Sir Vincent Wigglesworth (Wigglesworth, 1933; Wigglesworth, 1990). In 1933, he described cuticulin as an “amber coloured material, more or less mixed with melanin” and “a fatty and waxy substance” that he refined almost 60 years later as “a compost of sclerotin (quinone-tanned protein) with lipid”. Presence of cuticulin is indicated when after removal of wax, sclerotin interacts with silver ions in the argentaffin reaction in histological experiments. The molecular nature of cuticulin is, despite the long period of research, still enigmatic.

A major site of cuticulin accumulation is the outermost cuticle layer envelope. In *Drosophila melanogaster*, the envelope is constructed in three steps (Moussian et al., 2006a; Moussian, 2010). At the ultrastructural level, first, fragments of the envelope precursor consisting of two electron-dense sheets separated by an electron-lucid one are deposited into the extracellular space at the tips of irregular protrusions of the apical plasma membrane. Subsequently, these structures fuse to form a single layer covering the body of the developing embryo. Interestingly, fusion of envelope fragments at cell-cell contacts is the last step in this process (Fristrom and Liebrich, 1986). Next, when cuticle differentiation proceeds and the inner cuticle layers i.e. the epicuticle and the procuticle are formed, an electron-dense sheet intercalates into the electron-lucid sheet. The time point of this step of envelope maturation conceptually coincides with the time point of cuticulin deposition at the end of moulting in the kissing bug *Rhodnius prolixus* as observed by Wigglesworth (Wigglesworth, 1946). Transport and deposition of envelope material through the cuticle to the surface occurs via so called pore and wax canals, which are membranous tubes running from the apical surface of the epidermal cells to the surface of the cuticle. The underlying molecular mechanisms of lipid transport and integration into the cuticle or cuticulin formation are not understood.

In order to learn more about the mechanisms of lipid-based barrier construction, we conducted a screen for cuticle-deficient phenotypes caused by RNA interference (RNAi) in the late embryonic epidermis of *Drosophila melanogaster* that is mainly occupied with cuticle differentiation. Among others, we identified the ABC transporter CG9990 (Snustorr, Snu) and the extracellular protein CG2837 (Snustorr snarlik, Snsl) as essential factors involved in construction of the barrier against dehydration and penetration of water and xenobiotics. Based on our phenotypic analyses of *snu* and *snsl* we propose that Snu is involved in deposition of a yet uncharacterised, presumably lipidic material within the cuticle where it is immobilised by Snsl, which thus may constitute one of the protein components of Wigglesworth’s cuticulin.

## Results

In order to identify new factors of cuticle formation, we conducted a screen for cuticle defective phenotypes caused by RNAi-driven knockdown of transcripts that according to the modeENCODE and FlyExpress databases (Graveley et al., 2011; Kumar et al., 2011) are expressed in the late embryonic epidermis (see materials & methods).

### CG9990 and CG2837 are essential for integument barrier construction

Living larvae systemically expressing hairpin RNA against *CG9990* or *CG2837* mRNA, are more translucid than wild-type larvae (Fig. 1). When fixed with Hoyer’s medium, these larvae shrink, without folding or wrinkling of the cuticle (Fig. 1). *CG2837* knockdown larvae eventually hatch, but are flaccid and die around 10 minutes thereafter, whereas most and *CG9990* knockdown larvae fail to hatch. The tracheal system of these larvae is not gas-filled (Wang et al., 2015). Addition of halocarbon oil to newly hatched *CG2837* knockdown larvae rescues immediate lethality (Table 1). When manually freed from the egg case, larvae with reduced CG9990 function are unable to maintain their body shape, flatten and dry out.

**Figure 1.**
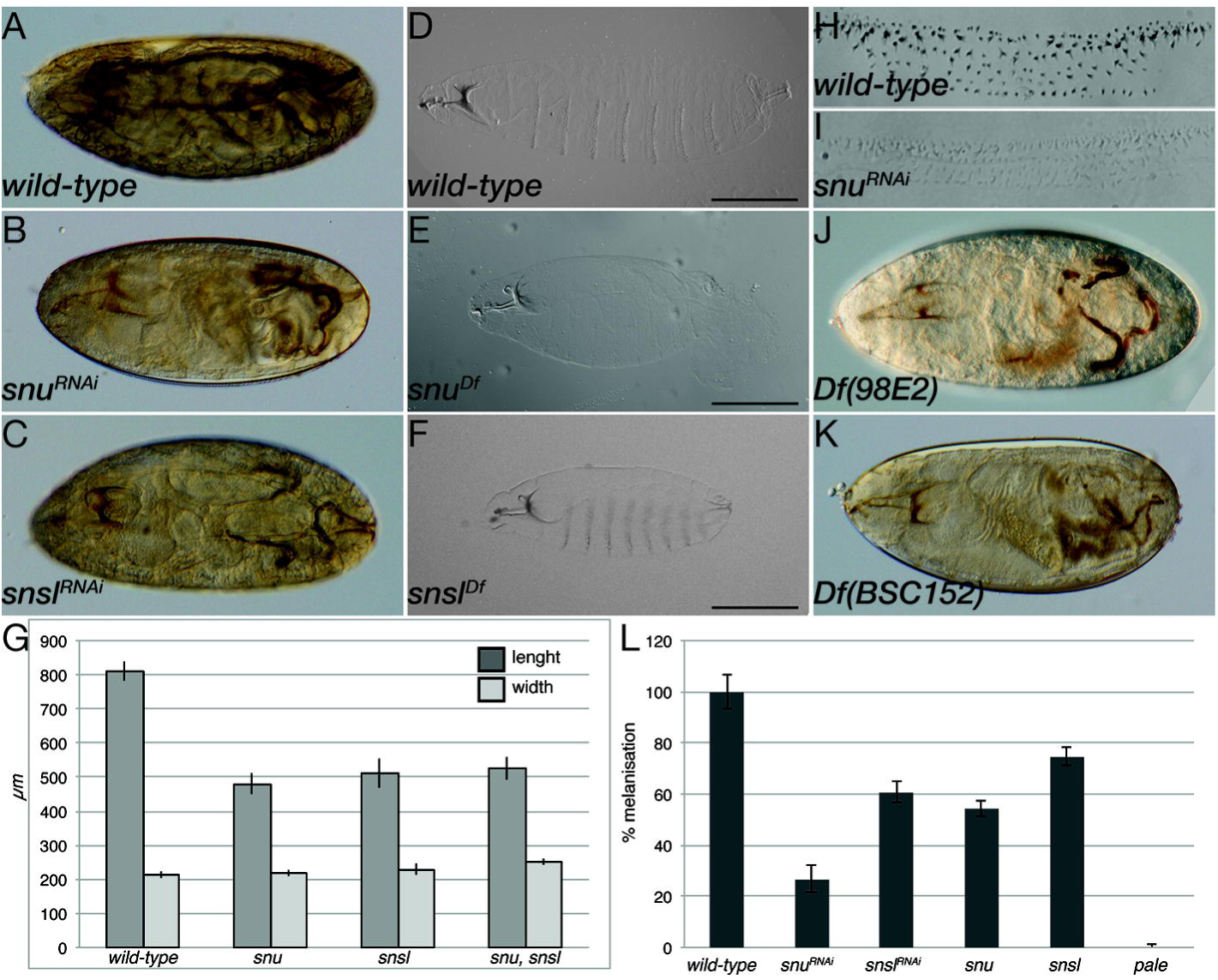
Phenotype of larvae with reduced or eliminated *CG9990/snu* and *CG2837/snsl* expression. At the end of embryogenesis, the larva ready to hatch is opaque, the inner organs are faintly visible; its tracheal system is filled with gas (A). Reduction of *snu* or *snsl* expression provokes a fully transparent appearance. The inner organs of these animals are clearly visible. Their tracheal system is not filled with gas (B,C). Cuticle preparations of wild-type (D) and *snu^Df^* (E) or *snsl^Df^* (F) mutant larvae reveal that the cuticle of *snu* and *snsl* larvae retracts. By consequence, the larvae are shorter (G). In particular in larvae with reduced *snu* expression, the head skeleton and the ventral denticles are paler (H,I). Live *snu^Df^* (J) or *snsl^Df^* (K) mutant larvae resemble those with reduced *snu* or *snsl* expression. Full melanisation of the head skeleton requires Snu, Snsl and Pale (L). Anterior is to the left, dorsal is to the top in A-F, J and K.

**Table 1.** Snsl prevents desiccation. In a simple desiccation test, wild-type and *snsl* knock-down larvae after hatching were either submerged in halocarbon oil or not. On air, *snsl* knock-down larvae died within 12 minutes, while wild-type larvae were alive during the 2 hours of experiment. Submersion of *snsl* knock-down larvae prolonged lifetime.

|  | wild-type | L370-Gal4xUAS*snsI*<sup>RNAi</sup> |
| --- | --- | --- |
| without oil | >2h (n=13) | 12min +/- 1.9 (n=10) |
| with oil | >2h (n=11) | >2h (n=11) |

To analyse the effects of complete *snu* elimination, we characterised embryos and larvae homozygous mutant for the embryonic lethal deletion *Df(98E2)* that removes *CG9990* and the neighbouring gene *huntingtin (HTT)*, which is dispensable for embryo development (Zhang et al., 2009). Indeed, expression of transgenic *CG9990* normalises the embryonic and larval *snu* mutant phenotype. Larvae homozygous mutant for *CG9990^Df(98E2)^* are translucid and rapidly dehydrate, when manually freed from the egg case, overall paralleling the RNAi-induced phenotype (Fig. 1).

To obtain a genetic stable constellation copying the *snsl* phenotype, we studied the embryonic phenotype caused by the deletion Df(2L)BSC182 that removes *CG2837* and six neighbouring loci. Overall, phenotypes provoked by Df(2L)BSC182 or systemic *CG2837^RNAi^* (daGal4/7063Gal4>UAS*CG2837^RNAi^*) are indistinguishable (Fig. 1).

These results together suggest that *CG9990*- and *CG2837*-knockdown larvae suffer rapid water loss. The desiccation phenotype prompted us to name *CG9990 snustorr* (*snu*, Swedish for bone-dry), *CG2837 snustorr snarlik* (*snsl*, Swedish for snu-like).

### Snu and Snsl contribute to the barrier against penetration

The cuticle is a bidirectional barrier, i.e. it does not only prevent water loss but also penetration of xenobiotics (Gibbs, 2011; Wang et al., 2016). To explore whether the cuticle of *snu* or *snsl* knockdown larvae is more permeable than the wild-type cuticle, we immersed these larvae in Eosin Y. Wild-type larvae remain unstained after incubation in Eosin Y at room temperature and at 40°C. They take up Eosin Y at 55°C. Reduction of Snu and Snsl function causes a dramatic uptake of Eosin Y through the integument already at 25°C or 40°C, respectively (Fig. 2). Applying this protocol, we also tested permeability of the wing cuticle in wild-type flies and flies expressing hpRNA against *snu* or *snsl* transcripts in the wing (Fig. 2). Wild-type wings do not take up Eosin Y at 25°C. At 50°C, by contrast, two areas in the posterior edge of the wings become red. Wings with reduced *snu* activity are impermeable to Eosin Y at 25°C, whereas the entire wing blade is stained by Eosin Y at 50°C. Reduction of *snsl* expression in wings does not interfere with Eosin Y uptake. Together, these findings demonstrate that *snu* and *snsl* are needed for inward barrier construction in the larval cuticle. In the wing cuticle, in contrast to Snu the function of Snsl seems to be dispensable. Thus, different types of cuticles have different requirements for Snu and Snsl.

**Figure 2.**
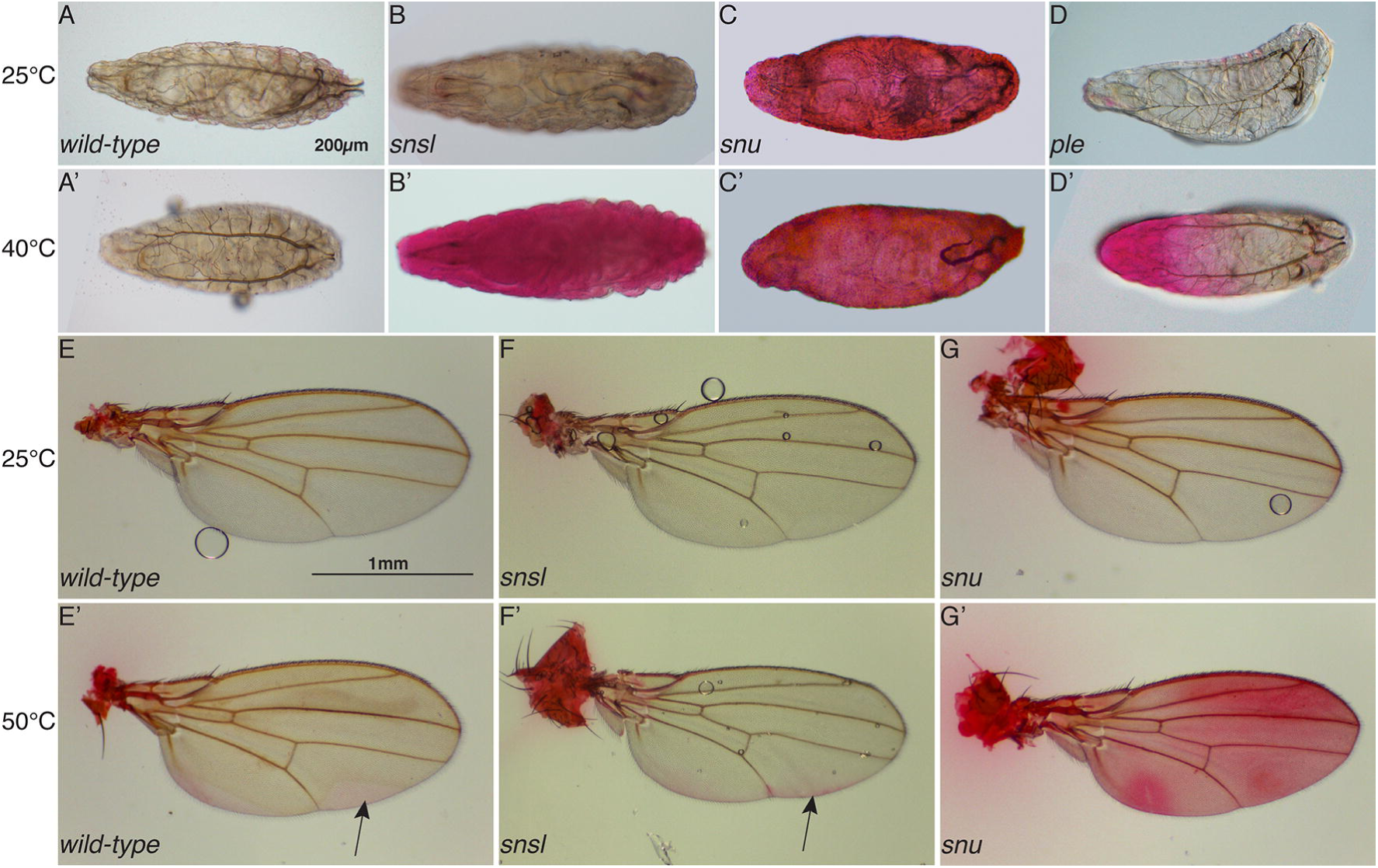
Snu and Snsl are needed for inward barrier function. The cuticle of wild-type larvae before hatching is impermeable to Eosin Y at 25°C and 40°C (A,A’). The cuticle of *snsl* mutant larvae becomes permeable to Eosin Y at 40°C (B,B’), whilst *snu* mutant larvae take up Eosin Y already at 25°C (C,C’). Larvae mutant for *pale* (*ple*) show Eosin Y leakage in the anterior body part at 40°C (D,D’). The wings of wild-type flies and flies with down-regulated *snsl* expression in the wings stain with Eosin Y at 50°C in the two areas of the posterior part (E-F’, arrows). The whole wings of the flies with down-regulated Snu activity in the wings are stained at 50°C (G,G’). A-D’ Dorsal view, anterior to the left. E-G’ Anterior to the top.

### Cuticle envelope structure requires Snu and Snsl function

The envelope, the outermost cuticle layer, is considered as the water impermeable barrier in insects. For a detailed analysis of Snu and Snsl function, we studied the cuticle ultrastructure of respective mutant larvae by transmission electron microscopy (TEM). The wild-type larval cuticle consists of three composite horizontal layers (Fig. 3) (Moussian, 2010). The outermost envelope has five alternating electron-dense and electron-lucid sublayers. Underneath lies the bipartite epicuticle. The innermost procuticle is a stack of helicoidally arranged sheets of parallel running chitin microfibrils (lamina). The envelope of *snu^Df(98E2)^* mutant larvae is reduced with only two electron-dense sublayers and occasionally breaks open. Unusual electron-dense material is found at the cuticle surface of these larvae. This material was observed by scanning electron microscopy (SEM), as well (Fig. 3). The epi- and procuticle, by contrast, appear to be normal in these larvae. In addition to the aberrant envelope and despite their normal ultrastructure, the procuticle detaches from the epicuticle in the cuticle at apodemes (muscle attachment sites) of *snu* deficient larvae (Fig. 3). The ultrastructure of the envelope of *snsl^BSC182^* mutant appears to be normal. However, unusual electron-dense material is present at the surface of these animals (Fig. 3).

**Figure 3.**
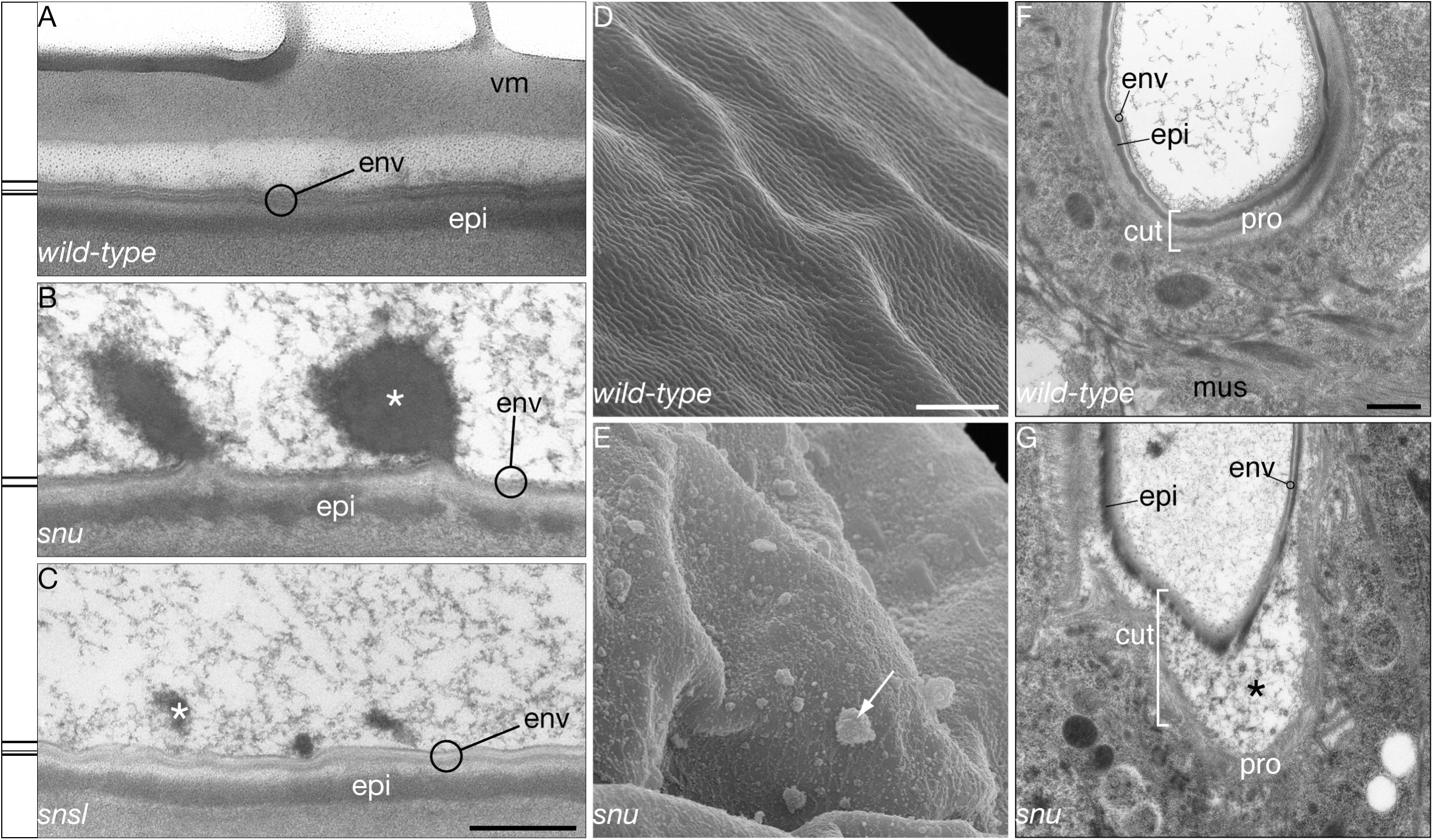
Snu and Snsl are needed for envelope construction. The envelope (env) of the wild-type first instar larva consists of three electron-dense and two electron-lucid alternating sheets above the epicuticle (epi) (A). In *snu* mutant larvae, the envelope is composed of two electron-dense sheets separated by one electron-lucid one (B). Electron-dense material protrudes from the surface of these animals (^∗^). The envelope of *snsl* mutant larvae appears to be normal (C). However, numerous electron-dense amorphic aggregates are found at the surface of these larvae (^∗^). The wild-type first instar larval surface forms minute ridges (D). By contrast, the surface of *snu* mutant first instar larvae is covered by irregular aggregates (arrow, E). At apodemes, the cuticle (cut) of the wild-type larva forms folds at muscle (mus) retraction (F). At these sites, the procuticle (pro) of *snu* mutant larvae detaches from the epicuticle (epi) when muscles retract (G). Scale bars: A-C shown in A 500nm; D and E shown in D 20μm, F and G shown in F 500nm.

In our experience, after excitation with 405nm violet light, the surface of the *D. melanogaster* L1 larva emits a distinct autofluorescent signal at a broad range of 420 to 620nm with the maximum peak around 475nm (Fig. 4). Further spectral analyses revealed a slight shift between the emission maxima at the upper and lower half of the fluorescing signal. This indicates that at least two materials with distinct optical properties form adjacent layers that constitute the surface of the *D. melanogaster* larva. At later larval stages additional dot-like structures (hereafter named 405-dots) are detected underneath the surface autofluorescence. These dots localise to the tips of filigree membrane protrusions, probably pore canals, visualised by CD8-RFP (Figs. 4, S1).

**Figure 4.**
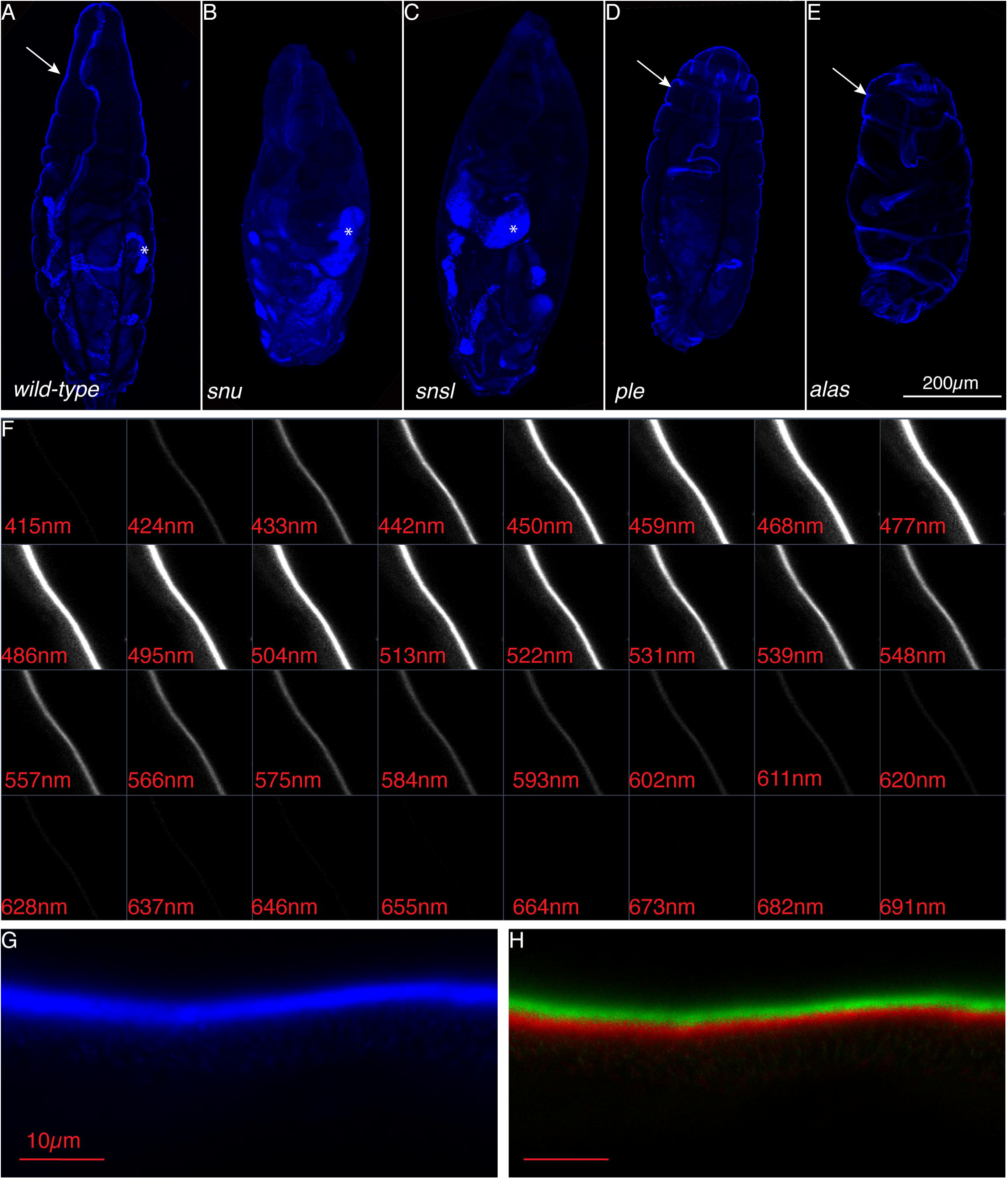
The larval envelope that fluoresces in a wide range of wavelengths is depleted in *snu* and *snsl* mutant larvae. When exposed to 405nm light, the cuticle of the WT *Drosophila* larvae before hatching emits a fluorescent signal (A). The cuticle of weakly melanised, permeable *snsl* and *snu* mutants does not emit this autofluorescent signal (B,C). Signal emission is normal in the cuticles of unmelanized *ple* (D) and *alas* mutants (E). Analysis of the emission spectrum of the surface autofluorescent light (lambda mode) of the cuticle of third instar larvae shows a broad range of emission between the boundaries of 424 and 620nm (F). The linear unmixing function reveals two potentially different substances in the autofluorescent region (G): the upper one is marked by green, and the lower part is marked by red (H).

When excited with this light, the surfaces of living first instar larvae suffering *snu* or *snsl* elimination display only a very faint autofluorescence. Thus, presence of the autofluorescing layer in *D. melanogaster* larvae depends on Snu and Snsl function. Our ultrastructural and fluorescence microscopy analyses together indicate that the surface autofluorescence depends on an intact envelope. Overall, we conclude that Snu and Snsl are involved in envelope construction.

### Envelope structure does not depend on melanisation or on di-tyrosine formation

In *snu^Df(98E2)^* larvae, denticles and the head skeleton are barely melanised (Fig. 1). To tackle the question whether the function of Snu might depend on melanisation, we analysed 405nm-excited surface auto-fluorescence of larvae mutant for *pale* (*ple^1^*) that codes for the first enzyme of the melanisation pathway. The surface auto-fluorescence of these larvae is not reduced compared to the signal in wild-type larvae (Fig. 4). This finding suggests that melanisation is not needed for envelope integrity. This is consistent with the finding that the envelope ultrastructure of embryos mutant for *Ddc* encoding the second enzyme of the melanisation pathway is normal (Moussian et al., 2006a). However, permeability experiments showed that the cuticle of *ple* mutants is permeable to Eosin Y at a level comparable to the cuticle of *snsl* mutant embryos (Fig. 2). This shows that either the disorders of the *ple^1^* envelope are not detectable by confocal or electron microscopy, or there are two parallel mechanisms implementing cuticle impermeability.

Previously, by analysing the phenotype of *δ*-*aminolevulinate synthase* (*alas*) mutant larvae, we had shown that a heme-dependent pathway is needed to prevent sudden water loss in the *D. melanogaster* first instar larva probably by introducing di-tyrosine bounds int the cuticle (Shaik et al., 2012). We had reported that the envelope of *alas* deficient 1^st^ instar larvae looked normal. Here, we tested, however, whether the surface autofluorescence in *alas* mutant larvae is intact (Fig. 4). When excited with a 405nm laser light source, the surface autofluorescence of these larvae appears to be normal. Thus, the heme biosynthesis pathway does not seem to contribute to the formation of the envelope as a waterproof barrier.

### Snu and Snsl are needed for cuticulin formation

According to Sir V. Wigglesworth the catecholamine-protein-lipid complex cuticulin contributes to cuticle waterproofness, tanning and stiffness (Wigglesworth, 1933; Wigglesworth, 1990). Concerning the pale cuticle, the desiccation problem and the ultrastructural defects of the envelope, it is well possible that cuticulin integrity is affected in *snu* and *snsl* mutant larvae. Cuticulin, the molecular nature of which is yet unknown, can be visualised by the so-called argentaffin staining (see materials and methods). In wild-type larvae, silver grains precipitate at the surface of the cuticle (Fig. 5). In *snu^Df(98E2)^* mutant larvae and larvae with reduced Snsl function, visibly less silver precipitates at the surface.

**Figure 5.**
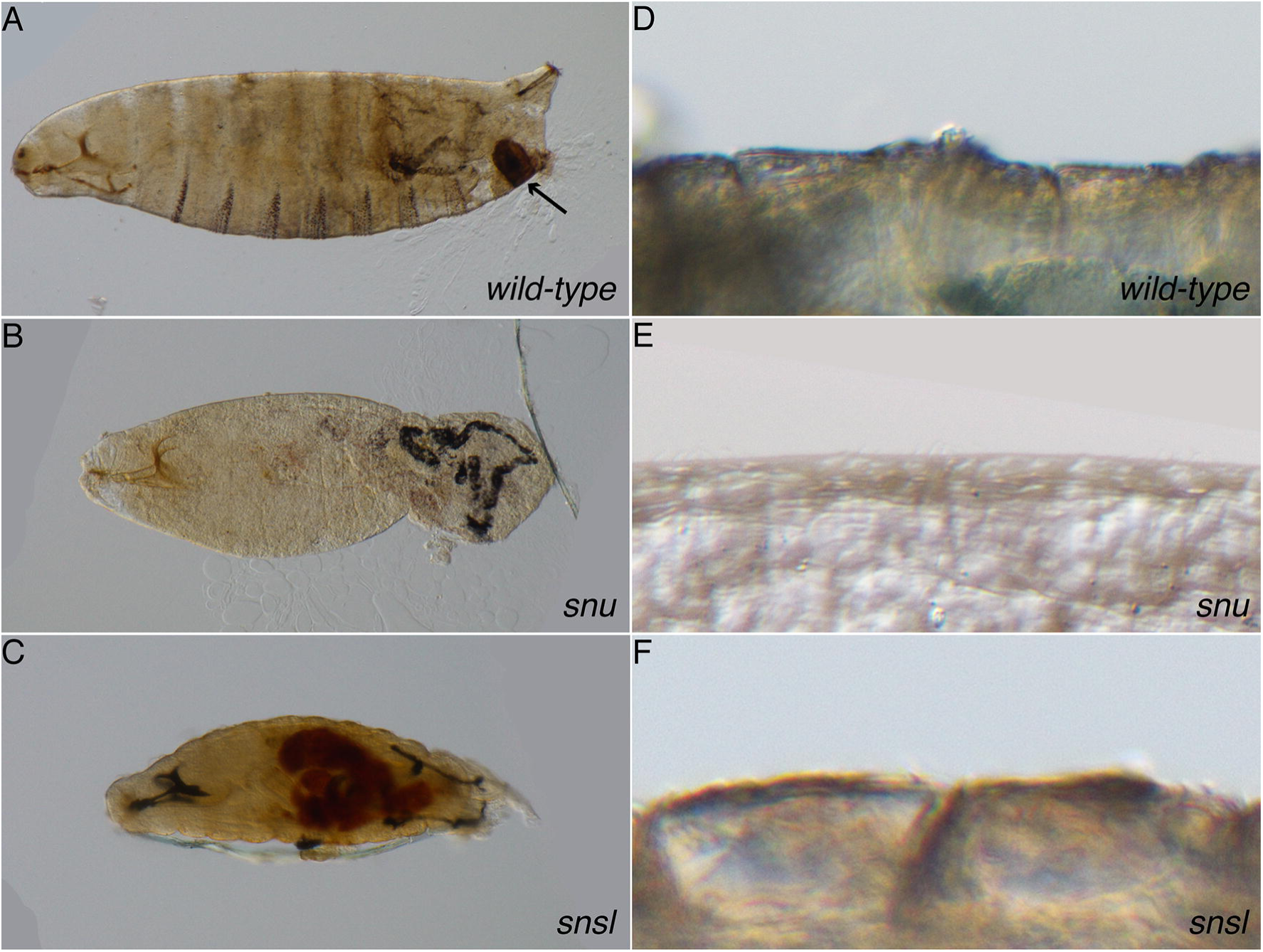
Cuticulin deposition depends on Snu and Snsl. Silver grains precipitate predominantly at the surface of the anal pads of wild-type first instar larvae following our argentaffin staining protocol based on Wiggelsworth’s protocol (A). Silver grains fail to recognise the surface of the anal pads of *snu* (B) or *snsl* (C) mutant first instar larvae. Sudan black marks the surface of wild-type first instar larvae (D). The surface of *snu* mutant first instar larvae is not stained by Sudan black (E), whilst the surface of respective *snsl* mutant larvae is (F).

We also sought to visualise cuticular lipids using Sudan black staining followed by light microscopy (Fig. 5). In wild-type and *snsl* mutant ready-to-hatch larvae, Sudan black stains the integument. By contrast, the integument of *snu* mutant larvae is unstained after incubation with Sudan black. Thus, in wild-type and *snsl* mutant larvae, integument lipids are present, while the amounts of lipids are reduced in the integument of *snu* mutant larvae.

Together, we conclude that Snu and Snsl are needed for cuticulin production.

### Snu codes for ABC transporters localised at the apical plasma membrane

To study the molecular and cellular function of Snu, we first analysed its protein sequence. The *snu* locus codes for six predicted isoforms of a half-type ABC transporter, the longest one, isoform A, harbouring 808 amino acids (Fig. S2). Standard homology searches reveal that all isoforms contain an ATP binding domain at the N-terminal half and a membrane-spanning region containing six transmembrane helices within the C-terminal half of the protein (flybase.org). The C-terminus is predicted to have an ER retention signal that upon interaction with a putative partner may be masked allowing the protein to continue travelling through the secretory pathway. Interestingly, none of the isoforms has a canonical N-terminal signal peptide. Snu-related sequences with the same domain composition are present in most arthropods suggesting that Snu function is conserved in this taxon (Fig. S2).

To verify the subcellular localisation of Snu, we generated an N-terminal GFP tagged version of Snu isoform A (GFP-SnuA) that is able to normalise the *snu* mutant phenotype (Fig. S3 and S4). At larval stages GFP-SnuA localises to the apical surface of epidermal cells of living larvae. Some GFP-SnuA dots are detected within the cell, possibly marking transporting vesicles. Hence, Snu is an ABC transporter that exerts its role in the cell and/or vesicle membrane.

### Snsl localises to the tips of pore canals

What is the molecular and cellular role of Snsl? According to our sequence analyses, Snsl is a secreted protein with an N-terminal signal peptide (Fig. S5). To verify the subcellular localisation of the Snsl protein, we generated a C-terminal RFP-tagged version of Snsl that is expressed under the control of an epidermal promoter (*twdlM*, see materials and methods). Expression of Snsl-RFP protein in *snsl* deficient embryos restores the autofluorescent surface layer excited with a 405 nm laser and cuticular impermeability (Fig. S4). The signal of Snsl-RFP is rather weak at early larval stages (Fig. 6), but well discernable at the third larval stage. In third instar larvae, Snsl-RFP is present in the cytoplasm and the cuticle. Here, it co-localises with the 405-dots underneath the envelope (Fig. 6). We assume that the dots composed of Snsl-RFP and the blue fluorescing material underneath the envelope are the tips of the pore canals running from the cell surface to the surface of the cuticle. To test this assumption, we analysed the localisation of these dots in third instar larvae expressing the membrane-inserted protein CD8-GFP that like CD8-RFP, albeit weaker, marks thin vertical lines in the cuticle presumably representing the pore canals. Snsl-RFP and the 405-dots contact the tips of CD8-GFP marked vertical lines supporting the notion that the Snsl-RFP/405-dots are at the ends of pore canals.

**Figure 6.**
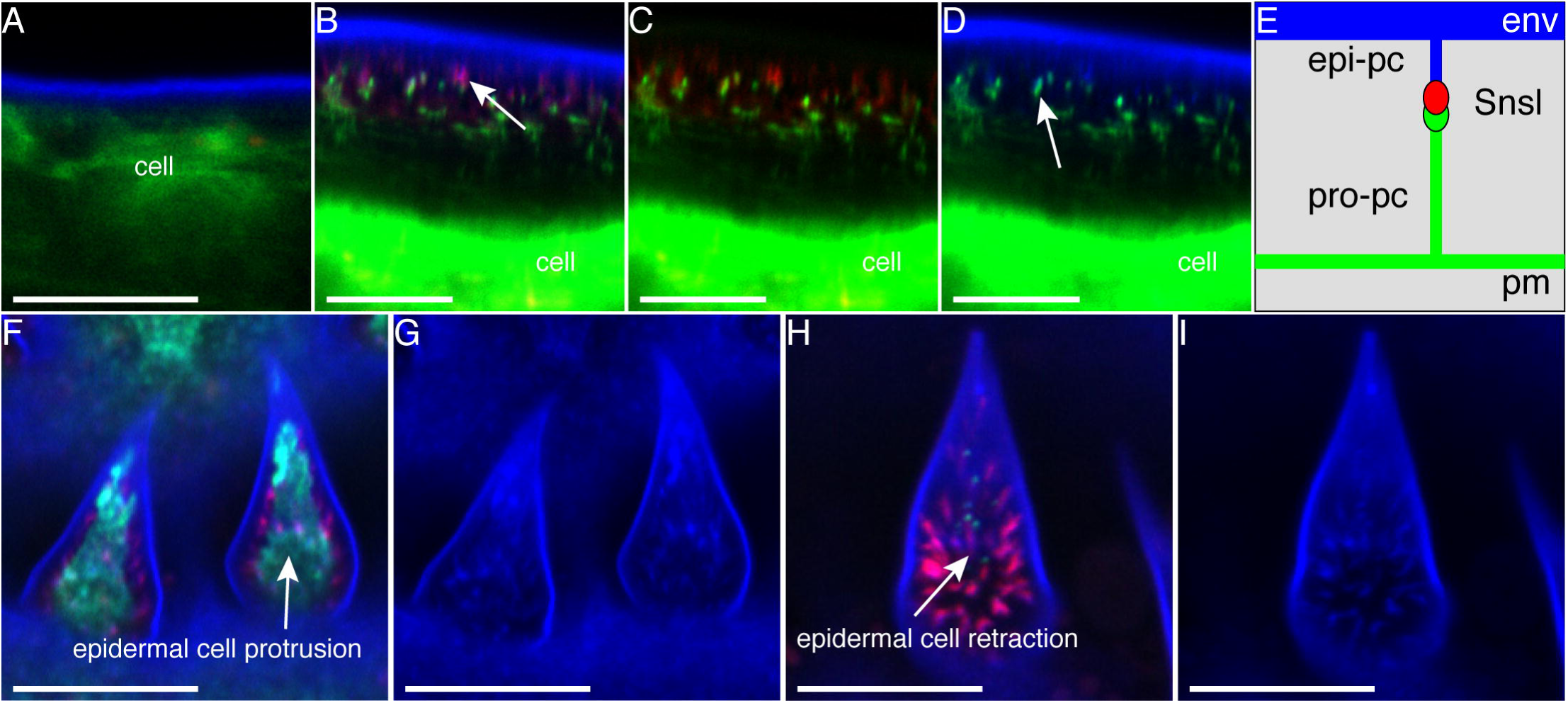
Snsl localises to pore canals. In first instar larvae, the Snsl-RFP protein (red) is visible as single dots in the epidermal cells that are marked in green. No red signal is detectable in the cuticle of first instar larvae (A) as in control larva not expressing Snsl-RFP (B). Snsl-RFP is visible in the cuticle of late third instar larvae, where it overlaps with the 405-dots and the tips of filigree membrane protrusions marked with CD8-GFP (C-D). Schematically, the epicuticular pore canals (epi-pc) are connected to the procuticular pore canals (pro-pc) through Snsl (E). These structures are visualised clearer by the expression of CD8-RFP in the epidermis (F). The epidermis (CD8-GFP, green) is present also in the middle of cuticular hairs of young third instar larvae (G) and retracts with time (H), when the distance between the cuticular envelope and the apical surface of epidermal cells increases. Snsl-RFP co-localises with autofluorescing (blue) canals of hairs that appear to connect the green epidermal cell with the envelope. Scale bar: 10μm.

### Localisation of Snsl in the cuticle depends on cell-autonomous Snu activity

The similar cuticle permeability problems associated with eliminated or reduced Snu and Snsl function argue that these proteins might collaborate or even interact with each other during envelope formation. Arguing that collaboration or interaction in the same process may necessitate co-localisation, we first tested whether the cytoplasmic localisations of Snu and Snsl coincide in live larvae. We observed that GFP-Snu and Snsl-RFP co-localise in the same dot-like structures within the epidermal cell, probably representing secretory vesicles (Fig. 7). These structures also contain fluorescing material when excited with the 405nm laser. The 405-dots and Snsl-RFP are found in the same structures when Snu expression is reduced by RNAi. Thus, GFP-Snu, Snsl-RFP and the 405 material are present in the same cytoplasmic structures. The presence of the 405 material and Snsl-RFP in these vesicles is independent of Snu function.

**Figure 7.**
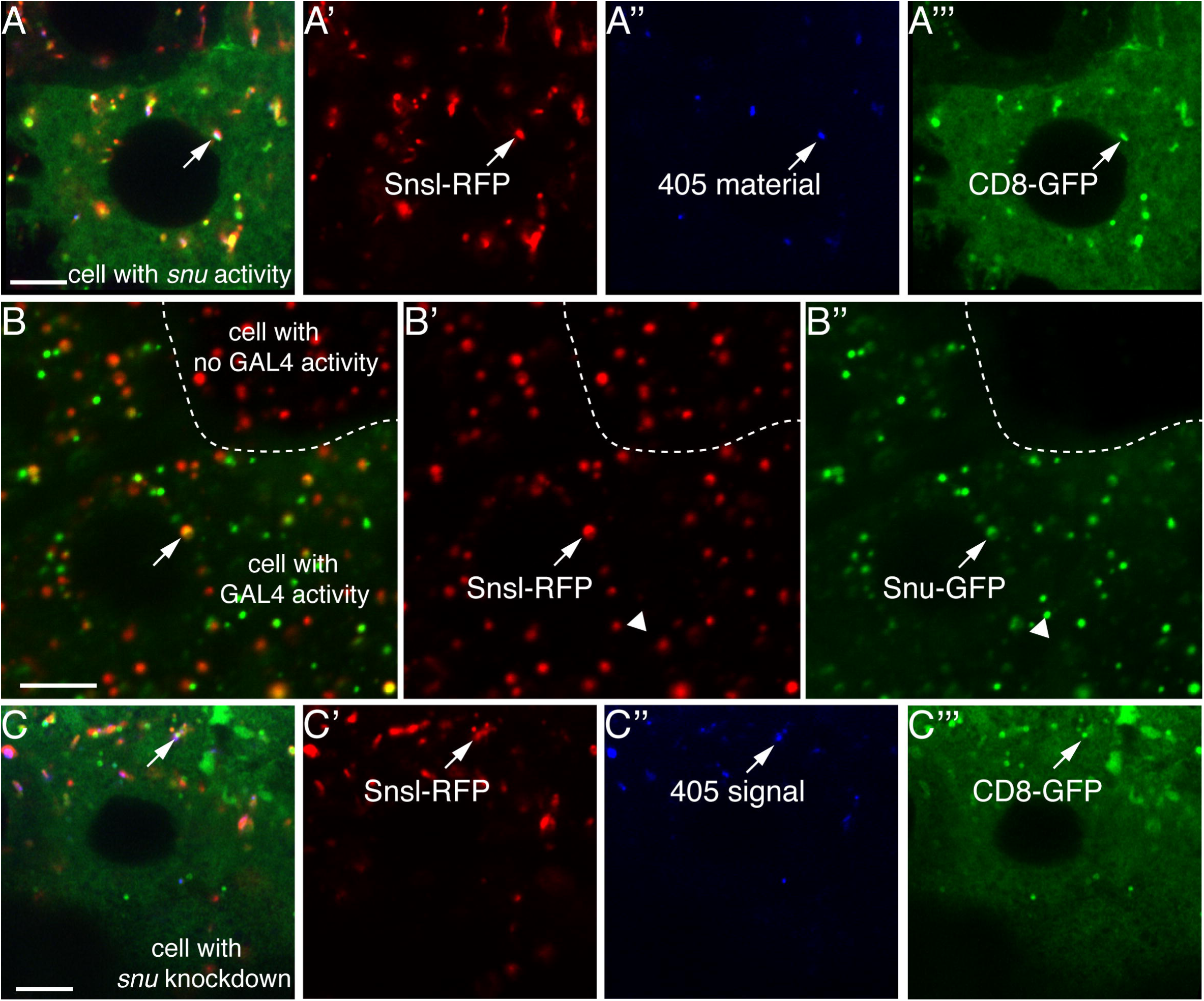
Co-localisation of GFP-Snu, Snsl-RFP and 405-dots precursors in secretory vesicles. In the epidermal cells of wild-type second instar moulting larvae, Snsl-RFP (red) and the 405-dot precursors (blue) are detected in the same structures (A-A’’’). These structures are also marked by membranous CD8-GFP (green) indicating they are transport vesicles. In the epidermal cells of wild-type second instar moulting larvae, Snsl-RFP (red) co-localises with GFP-Snu (green) in the same vesicles suggesting that these two proteins may interact (B-B’’). The white triangle points to a Snu-GFP-vesicle that is devoid of Snsl-RFP. The Gal4 driver used in this experiment has a mosaic expression. In the cells with down-regulated *snu* expression (C-C’’’) 405-dots and red Snsl-RFP still co-localise in the membranous structures indicating that Snu is not required for the vesicle loading process.

We next tested whether the extracellular localisation of Snsl-RFP depends on Snu function in larvae mosaic for Snu activity (Fig. 8). In cell clones with reduced Snu function, Snsl-RFP localises in the presumed procuticle. The Snsl-RFP signal in these cells overlaps with the ectopic 405-signal. Thus, correct delivery of Snsl and of the 405-dots is dependent on full Snu function.

**Figure 8.**
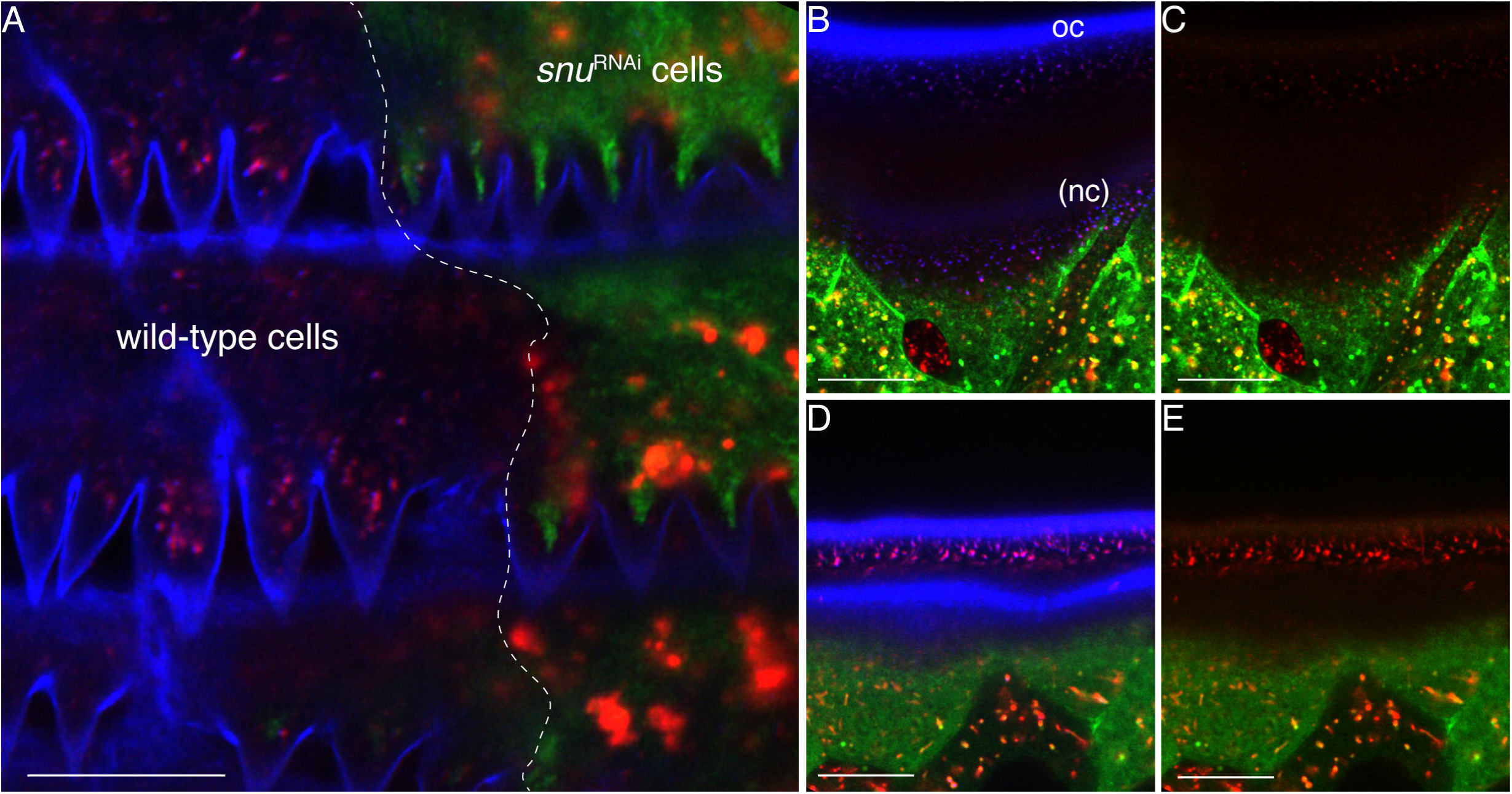
Snsl localisation depends on Snu function. As observed in tangential optical sections, in the clones with RNAi-induced *snu* expression reduction (marked with GFP, green) the autofluorescence signal (blue) is weaker (A). The Snsl-RFP signal (red), which overlaps with the 405-dots in the pore canals in the extracellular space in wild-type clones, is intracellular in clones with reduced *snu* expression. In sagittal optical sections at higher magnification, below the very faint envelope of newly formed *snu*-deficient third instar larval cuticle (nc), 405-dots accumulate in the lower cuticular layers above the epidermal cells and colocalised with the Snsl-RFP signal that is discernable at this magnification (B,C). The old cuticle (oc) established before the onset of RNAi is intact. In the extracellular space of respective control larvae accumulation of 405-dots and Snsl-RFP signal does not occur (D,E). Scale bars: 10μm

## Discussion

Sir V. Wigglesworth proposed in 1933 that the catecholamine-lipid-protein complex cuticulin is a key factor in cuticular transpiration control in *Rhodnius prolixus* (Wigglesworth, 1933; Wigglesworth, 1990). After more than 80 years, the molecular composition of cuticulin is, however, still elusive. In the present work, we report on the identification of three factors that we propose to be involved in cuticulin formation in the fruit fly *Drosophila melanogaster.*

### Course of envelope construction

The cuticle envelope is a thin composite structure. It is composed of five alternating electron dense and electron lucid sheets (Moussian et al., 2006a). The *D. melanogaster* larval envelope is produced and assembled during embryogenesis sequentially (Moussian et al., 2006a). Fragments of envelope precursors with two electron-dense layers framing an electron-lucid layer are deposited into the extracellular space. They eventually fuse together forming a continuous sheet. An electron-dense layer intercalates within the electron-lucid layer and the envelope maturates. Extensive protrusions of the apical plasma membrane called pore canals are reported to be the routes of surface i.e. envelope material delivery (Locke, 1961). In this work, we show that the surface of *D. melanogaster* larvae fluoresces when excited with a 405 nm light source. During cuticle formation, dots fluorescing upon illumination with this light source are detected within the epidermal cell and in the forming cuticle. Some of these dots also localise to the tips of the pore canals (405-dots). We reckon that 405-dots are components that are transported via the pore canals and are incorporated at the surface of the larvae constituting the envelope. Persistent localisation of this signal at the tips of pore canals also indicates that a second population of this material additionally represents constituents of the pore canals. The relationship of the 405-dots and the ultrastructure of the envelope and pore canals remains to be investigated.

### Snu and Snsl are needed for envelope construction and function

Reduction or elimination of *snu* expression results in rapid water loss, uncontrolled uptake of external dyes and death of the larvae. As shown by electron microscopy, these larvae have a thin and depleted envelope. Thus, Snu is not needed for the first step of envelope formation. Depletion of the envelope is also recognisable with the fluorescence microscope. The surface, 405nm-excited fluorescence observed in wild-type larvae is lost in animals with reduced or eliminated Snu function. Instead, dotted 405 signal is detected within the procuticle and occasionally in the cell in these animals. At the ultrastructural level, we observe that an electron-dense sublayer, possibly the intercalated one, is missing. Furthermore, the integument of *snu* mutant larvae remains unstained after incubation with the lipid-marking dye Sudan black. Additionally, in argentaffin staining assays, the surface of *snu* mutant larvae fails to be marked with silver grains. These results indicate that lipophilic components in or at the cuticle are depleted when Snu is dysfunctional. The putative orthologues of Snu in *Tribolium castaneum*, TcABCH-9C, and *Locusta migratoria*, Lm TcABCH-9C have consistently been reported to be needed for the presence of lipids at the surface of the larvae, which presumably are in turn required for desiccation resistance (Broehan et al., 2013; Yu et al., 2017). We conclude that the ABC transporter Snu is needed to establish the envelope as a water-resistance barrier by mediating the incorporation of the 405-dots and lipophilic components into this layer.

A similar, albeit weaker larval phenotype is caused by reduction or elimination of *snsl* expression. These larvae are able to hatch after embryogenesis and die as first instar larvae that massively lose water. Dye penetration into these larvae occurs at higher temperatures than into *snu* deficient larvae, the cuticle of wings with reduced *snsl* expression remains impermeable to Eosin Y, and Sudan black detection of lipophilic components in the integument is positive. Attenuated dye penetration in *snsl* deficient larvae may be due to presence of these Sudan black-sensitive lipophilic molecules on their surface and their absence on the surface of *snu* deficient larvae. In any case, similarly to the *snu* mutant phenotype, based on ultrastructural analyses and by fluorescence microscopy, the envelope of larvae with reduced or eliminated Snsl function appears to be aberrant or depleted. Based on the localisation of the Snsl protein at the tips of pore canals, we conclude that it indirectly participates in the construction of the envelope as an impermeability barrier. Moreover, the similar phenotype of larvae with reduced or eliminated *snu* and *snsl* function suggests that the respective proteins act in the same mechanism of envelope formation and function.

### Snu and Snsl function is independent of the heme-biosynthesis pathway

Recently, we discovered that the heme-biosynthesis pathway is needed for water retention by the *Drosophila* larval cuticle (Shaik et al., 2012). A dityrosine network produced by a yet unknown heme-dependent peroxidase probably within the procuticle is reduced in these animals. Their envelope is, by contrast, normal and impermeable. Thus, the barrier formed in dependence of the heme biosynthesis pathway and the one formed by the presumed Snu-Snsl pathway are separate structures. The Snu-Snsl-dependent envelope-borne barrier does not involve a di-tyrosine network.

### Snu and Snsl cooperate during envelope construction

How may Snu and Snsl act together in envelope construction and function (Fig. 9)? Snu is an ABC transporter that localises to vesicles and the apical plasma membrane of epidermal and tracheal cells during cuticle formation. It is required for correct localisation of the 405-dots and the extracellular protein Snsl-RFP in the mature cuticle. The Snu-positive vesicles also contain Snsl-RFP and the 405-dot material. In *snu* deficient animals, Snsl-RFP and the 405-dots co-localise in vesicles suggesting that loading or formation of these vesicles or membrane structures with 405-dot material and Snsl-RFP is independent of Snu function. By contrast, Snsl-RFP and the 405-dots are mislocalised in the cuticle of *snu* deficient larvae. This finding indicates that the content of the 405-dot and Snsl-RFP vesicles is not functional in the absence of Snu. Hence, in summary, Snu mediates loading of these vesicles with a substance that is needed for the successful localisation of the 405-dots and Snsl-RFP in the extracellular space after secretion. Conceivably, our lipid staining experiments with Sudan black argue that these components are lipophilic. This material, is present in *snsl* deficient (with normal Snu activity), but not in *snu* deficient larvae. In conclusion, the 405-dots, Snsl-RFP and the substrate of Snu together constitute an essential element of the envelope and of the sub-envelope region as a waterproof barrier.

**Figure 9.**
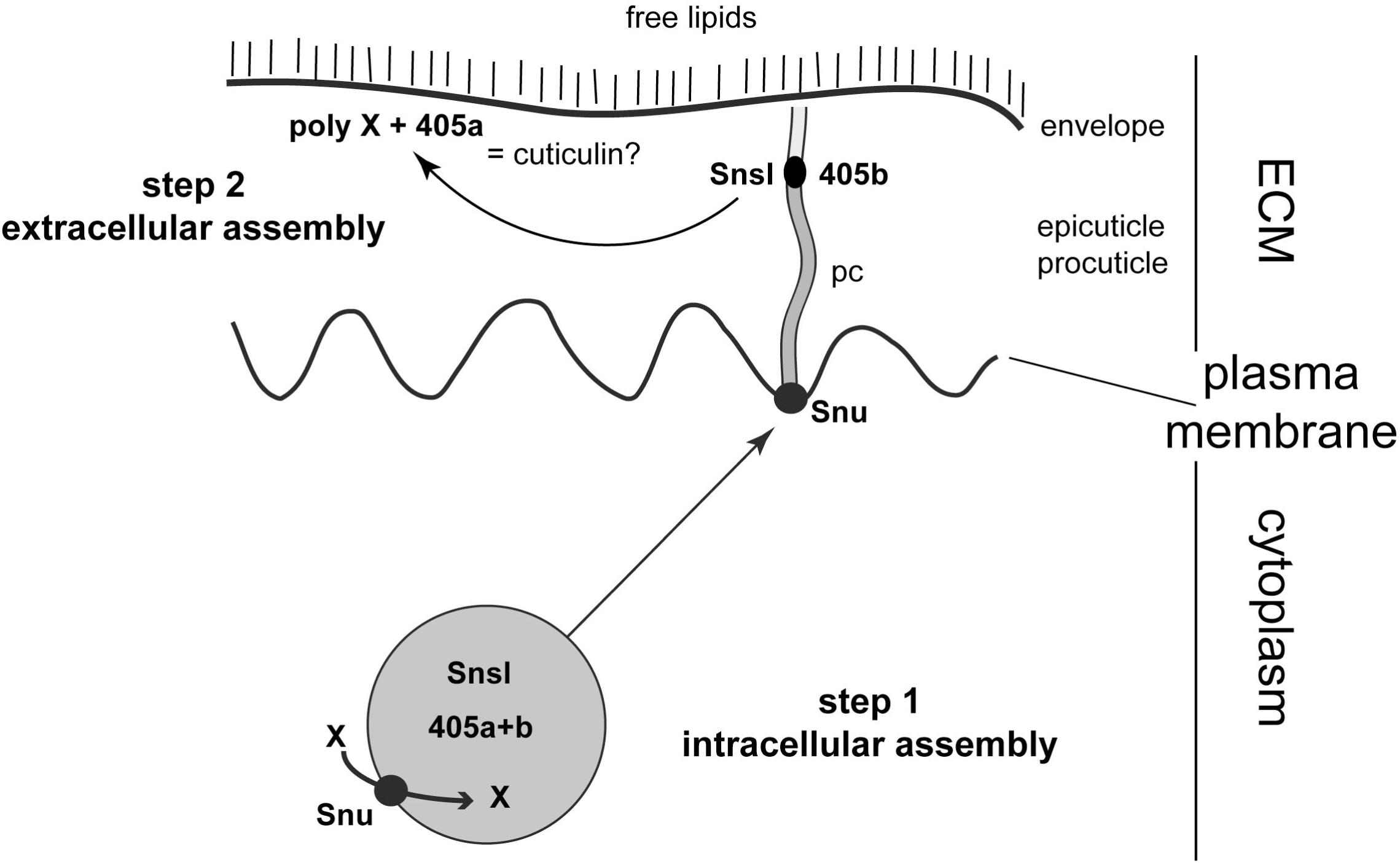
Model for Snu and Snsl function. Snu localises to vesicles and is probably needed to load cuticle material (X) into these vesicles that also contain Snsl and 405-dot precursors (405a+b). In these vesicles, a first step of inward barrier assembly may occur. They eventually fuse with the apical plasma membrane, and their content is released into the differentiating cuticle. Possibly via the pore canals, Snsl and a subfraction of the 405-dot precursors (405a) are delivered to the distal regions of pore canals. Another subfraction of 405-dot precursors (405b) and probably the Snu substrate X are used to form the envelope. The distribution of this subfraction of 405-dot depends also on Snsl function. This second step of assembly in the extracellular space establishes the inward barrier of the cuticle.

This scenario implies that localisation of Snu in the apical plasma membrane is the consequence of vesicle fusion with the plasma membrane and that it has no significance for Snu function per se. In an alternative scenario, Snu may play an important role in the apical plasma membrane. Following this line of argument, Snu may be involved in the formation of the membranous pore canals that are important routes of material transport to the cuticle surface including the envelope. Mislocalisation of Snsl-RFP and the 405-dots in *snu* deficient larvae would be a consequence of defective or inexistent pore canals. These alternative scenarios are shown in figure 9.

### Parallels to vertebrate lipid-based ECM formation

In vertebrates, the extracellular pulmonary surfactant that stabilises alveolar structure during breathing is produced by ABCA3 that localises to alveolar type 2 epithelial cells (Beers and Mulugeta, 2017). Precursors of the surfactant are assembled first in ABCA3-harbouring lamellar bodies that derive from multi-vesicular bodies. These vesicles are transported to the apical plasma membrane and their contents are released into the extracellular space. ABCA3 is eventually recycled back to lamellar bodies or degraded by lysosomes. Likewise, in keratinocytes, ABCA12 is essential for the formation of the lamellar bodies inside the cell before deposition into the extracellular space by a yet unknown mechanism (Feingold and Elias, 2014).

In *D. melanogaster*, a portion of Snu localises to intracellular vesicles that contain envelope precursors including lipids and Snsl, in addition, some other portion of Snu localises to the apical plasma membrane, while some Snu-positive vesicles within the cell, probably recycling endosomes, are devoid of Snsl and envelope precursors.

In general, thus, production of lipid-based extracellular matrices may by default involve the activity of an ABC transporter that is needed to assemble minimal matrix units within intracellular vesicles. Based on our previous work on the function of the t-SNARE Syntaxin-1A (Syx1A) showing a depleted envelope in *syx1A* mutant larvae (Moussian et al., 2007), we reckon that subsequent release of the ECM units to the extracellular space may indeed depend on Syx1A. This mechanism is elusive in vertebrates.

### Does the assembly of Wigglesworth’s cuticulin depend on Snu?

Cuticulin is the key component of the envelope as a waterproof barrier (Wigglesworth, 1970; Wigglesworth, 1975). It is a complex composed of “wax and sclerotin” (Wigglesworth, 1985a), which itself is a protein-quinone complex of unknown composition. To what extend are our findings conform to the classical cuticulin concept of Wigglesworth?

Several aspects of Wigglesworth’s cuticulin function are compromised in larvae with reduced or eliminated *snu* or *snsl* expression. Most importantly, the envelope of these animals is depleted and is unable to control inward and outward flow of water or xenobiotics (Eosin Y). Additionally, their head skeleton and denticles fail to melanise normally. Conceptually, following Wigglesworth’s arguments, cuticulin assembly or function appears to be disrupted in these animals. Both proteins, however, do not localise to the envelope, but to a sub-envelope region (Snsl) or the apical plasma membrane (Snu). These data, hence, suggest together that Snu and Snsl are not constituents of the envelope or cuticulin itself but mediate envelope construction possibly by assisting cuticulin assembly. The sub-envelope pore canal population of cuticulin i.e. 405-dots, on the contrary, do harbour Snsl that may play a role in sealing the pore canals.

A core component of cuticulin may be represented by the 405-material in the envelope and the 405-dots below the envelope. The 405-dots may be distinct structures or merely the storage or transport form of the 405-material. The absence of Snsl in the envelope and depletion of the envelope in *snsl* mutant larvae argue together that the 405-dots are transported to the envelope in an Snsl-dependent mechanism, a process that starts already within the cell. For a full comprehension of the relationship between cuticulin and the 405-material, Snu and Snsl the molecular composition of the 405-material needs to be identified and characterised.

## Methods

### Fly husbandry

For embryo and larva collection, flies were kept in cages on yeasted apple juice agar plates. Embryos were staged according to the gut morphology and the time of development at 25°C described in Hartenstein and Campos-Ortega (Hartenstein and Campos-Ortega, 1985) and dechorionated for three minutes in commercially available chlorine bleach diluted 1:1 in tab water. Homozygous or transheterozygous mutant embryos and larvae were manually collected in a population of progeny segregating a respective balancer chromosome expressing GFP (Dfd>YFP or Kr>GFP) and the mutation-bearing non-GFP chromosome. For techniques that require removal of the vitelline membrane the embryos were devitellinized by shaking in equal amounts of heptane and methanol (fixed embryos), or manually in PBS buffer (live embryos).

### RNAi screen and Snu and Snsl knockdown larvae

For the RNA interference (RNAi)-based screen for cuticle phenotypes, the Berkley *Drosophila* Genome Project (BDGP) *in situ* database was first searched for genes that are specifically expressed in the epidermis during mid to late stages of development when the cuticle is formed. Next, to analyse the function of a given factor during embryonic cuticle formation, flies harbouring the respective UAS-RNAi construct from the Vienna *Drosophila* RNAi Centre (VDRC) (Dietzl et al., 2007) were crossed to flies expressing the epidermal Gal4-driver 69B-*Gal4* and *dicer2* (UAS-*dicer2*) that has been reported to enhance RNAi efficiency. In total, 129 UAS-RNAi lines representing 75 candidate genes were tested. As a first indication for the importance of a factor for embryonic cuticle formation, embryonic lethality was assessed. When lethality of the offspring exceeded 30%, cuticle preparations of the lethal fraction were made following the protocol of Nüsslein-Volhard and Wieschaus (Nüsslein-Volhard et al., 1984) to analyse the phenotype. As controls for screen efficiency, RNAi silencing of *krotzkopf verkehrt* (*kkv*) and *knickkopf* (*knk*) (Moussian et al., 2005; Moussian et al., 2006b) was conducted. As a negative control flies harbouring the Gal4 driver were crossed to *w^1118^* flies.

For further phenotypic analyses of *snu* (#107544 VDRC) and *snsl* (#104558 VDRC), flies with the respective UAS-RNAi construct were crossed to flies expressing both Gal4-drivers *da*-Gal4 and *7063*-Gal4 (maternal Gal4). Crosses were generally kept on yeasted apple juice agar plates at 25^o^C over night. *snu* RNAi L2/L3 knockdown larvae were additionally obtained by crossing *snu* UAS-RNAi flies with *hsp70*-Gal4 (#2077, BDSC) flies. Gal4 expression by *hsp70*-Gal4 is leaky in some epidermal cells; hence, to avoid heat shocking that may damage larvae, we developed a protocol taking advantage of the Gal4 leakiness. Larvae were kept at 18°C until early L2 stage. Then they were transferred to 20°C, 22°C, 25°C and 28°C, respectively, to avoid heat-shock induction, and enhanced RNAi effect during L2/L3 transition in the leakage areas.

### Rescue and localisation experiments

Total mRNA was prepared from OrR wild-type stage 17 embryos (RNeasy Micro Kit, Qiagen). Total cDNA was prepared from the mRNA (Superscript III First-Strand Synthesis System, Invitrogen), which was used as a template for amplification of *snu* (Expand High Fidelity PCR System, Roche). Two alternative 5’ primers, ATGGCGCCGAAGAAAGAGG and ATGCTGATATCAACTATTTCCAC, and one 3’ primer, CCAAACGCGGCTAGGATCTT were used for the amplification of three different isoforms. The products were ligated into the pCR2.1 TOPO vector (Invitrogen), which was used as a template for another PCR reaction with the same primer combinations. The product was ligated into the pCR8/GW/TOPO vector (Invitrogen). Sequencing of the purified vector DNA (Miniprep kit, Qiagen) confirmed the presence of *snu* isoforms A and B. Next, LR recombination was used to clone the cDNA into Gateway vector pTGW, which contains a UAS promoter and a GFP tag N-terminal to the insert. The vector was amplified in DH5alpha *E. coli* bacteria, purified (PerfectPrep Endofree Maxi Kit, 5Prime) and sent to Fly-Facility, (Clermont-Ferrand, France) for injection into *D. melanogaster* embryos.

For rescue studies using the UAS-GFP-SnuA construct, the UAS-GFP-SnuA insertion on the third chromosome was recombined on the chromosome harbouring the *snu^Df(98E2)^* allele to obtain the UAS-GFP-SnuA, *snu^Df(98E2)^* chromosome. Similarily, the insertions *tub*-Gal4 or 69B-Gal4 were recombined to the *snu^Df(98E2)^* carrying chromosome to obtain the *tub*-Gal4/69B-Gal4, *snu^Df(98E2)^* chromosome. Flies with UAS-GFP-SnuA, *snu^Df(98E2)^* were crossed to fies with *tub*-Gal4/69B-Gal4, *snu^Df(98E2)^* in rescue experiments. To study Snu localisation, UAS-GFP-SnuA harbouring flies were crossed to *knk*-Gal4 or *da*-Gal4 flies and analysed by confocal microscopy.

In order to rescue the Snsl-deficient phenotype, flies with the deficiency uncovering the *snsl* gene Df(2L)BSC182 were crossed to flies with TwdlM>Snsl-RFP on the first chromosome. The TwdlM>Snsl-RFP construct was synthesised by Genewiz at Sigma-Aldrich.

### Histology, microscopy and image preparation

For live imaging, larvae were anesthetised with ether, mounted in glycerol or halocarbon 700 on a slide and covered by a coverslip. They were observed using Zeiss confocal microscopes (LSM 710, 780 or 880).

For the analysis of the emission of the autofluorescent substance, a Zeiss LSM 880 using the ZEN software was applied. Cuticle optical cross-section of live third instar *D. melanogaster* larvae was scanned with 405nm laser light (Supplementary fig.7). Emission light was collected at wavelengths ranging of 415-691nm in 8-9nm sub-ranges creating 36 pictures. On a collective lambda mode picture (composed of superimposed 36 pictures) small round-shaped areas were designated and for each area a graph showing intensity - emission wavelength dependence was generated. All graphs from all areas were compared and divided into two groups. Graphs of these two groups showed a small shift of the maximum intensity. Two of the areas representing these two different types of graphs were taken as a reference and applied in a “linear unmixing” mode. The whole autofluorescent signal was divided into two parts corresponding to one of the graph type.

In penetration assays, live larvae were incubated in a 0,5% Eosin Y staining solution (Sigma-Aldrich) for 20 minutes at 25°C and 40°C, and subsequently washed with tap water before mounting for microscopy with a Leica Z4 using an unbuilt camera and Leica software. Wings of flies generated by the cross of *vg*-Gal4 flies with UAS-*snu* or UAS-*snsl* flies were stained with Eosin Y following the same protocol. Bromophenol blue penetration was tested with 0,1% Bromophenol blue (Sigma-Aldrich) at room temperature for five minutes. Larvae were washed three times with tap water, and viewed on a Leica Z4 binocular.

For Sudan Black B staining, after dechorianation and mechanical removing of the vitelline membrane, wild-type, *snu* and *snsl* mutant stage 17 embryos were incubated with 100μl Sudan Black B solution (0.5% in 70% ethanol, Sigma-Aldrich) over night at room temperature. Embryos were washed three times with 70% ethanol and incubated in 70% for six hours for destaining. Specimens were transferred to a glas slide, embedded in halocarbon oil and analysed by light microscopy.

For argentaffin detection, we modified a protocol published by Wigglesworth (Wigglesworth, 1985b; Wigglesworth, 1985a) for staining *D. melanogaster.* Five to ten stage 17 embryos were freed manually from the egg case using needles after dechorionation in 50% bleach for 3 minutes. They were collected in 200 μl of 5% (w/v) AgNO_3_. After collection, 800 μl of 2.5% (w/v) Tris base were added. Larvae were incubated in this solution for one hour at 60°C. Staining reaction was stopped with 1ml of a 2.5% Thiosulphate solution. Stained larvae were analysed using a Nikon AZ100 microscope with a Nikon camera and NIS software.

For transmission electron microscopy (TEM), embryos and larvae were treated as described in (Moussian and Schwarz, 2010). A Philips CM-10 was used at 60kV for data recording. For scanning electron microscopy (SEM), we applied a protocol published in Moussian 2006.

Figures were prepared using the Adobe Photoshop CS3 and Adobe Illustrator CS4 software without changing the initial settings of the microscope.

## Acknowledgments

We thank FlyBase and the Bloomington Drosophila Stock Center, for information and fly stocks.

## Author contributions

Conceived and designed the experiments: MN, RZ and BM. Performed the experiments: MN, RZ, YW, KO, JB, NG, DA, MF and BM. Analyzed the data: MN, RZ, DA and BM. Wrote the paper: RZ and BM.

---

Supplementary data 1. Visualisation of the pore canals through the expression of UAS-CD8-RFP, which gives a stronger signal than UAS-CD8-GFP shown in figure 6.

Supplementary data 2. Multiple sequence alignment of Snu and putative Snu orthologues from different arthropod species using CLUSTAL O (1.2.1).

All orthologues belong to class H of ABC transporters. ATP-binding domain marked in blue (residues 74-286), transmembrane helices predicted by PSORTII (https://psort.hgc.jp/cgi-bin/runpsort.pl) marked in red (residues 433-802). All orthologues have an ER retention signal predicted by PSORTII at the very end of the sequence marked in bold (RAKR in Snu).

Supplementary data 3. Localisation of GFP-Snu does not depend on Snsl function.

GFP-SnuA (green) ubiquitously expressed localises to the apical plasma membrane of epidermal cells (a). Some signal is observable also inside the cells. The surface autofluorescence marks the envelope (blue). The localisation of GFP-SnuA in first instar larvae is similar (b), also in the animals with down-regulated Snsl activity (c). Note that in these larvae the surface autofluorescence is strongly reduced.

Supplementary data 4. Rescue of the *snu* and *snsl* phenotypes

Expression of GFP-SnuA protein driven ubiquitously restores the surface autofluorescence of the *snu* mutant embryos (1a-1c: wild-type, *snu* mutant and *snu* mutant containing GFP-SnuA, respectively). It also restores cuticle impermeability (3a-3a’ wild-type before and after Bromophenol blue application; 3b-3b’ *snu* mutant; 3c-3c’ *snu* mutant with GFP-SnuA). Expression of Snsl-RFP in the background of *snsl* mutant embryos restores the surface autofluorescence (2a-2a’ wild-type; 2b-2b’ *snsl* mutant embryo; 2c-2c’ *snsl* mutant embryo containing Snsl-RFP) and impermeability of the cuticle (3d-3d’ *snsl* mutant; 3e-3e’ *snsl* mutant with Snsl-RFP).

Supplementary data 5. Alignment of *D. melanogaster* protein sequence of Snsl (CG2837) with sequences of its putative orthologues from other insect species (*Bombyx mori*, *Tribolium castaneum* and *Apis mellifera*) using the CLUSTAL O (1.2.1) multiple sequence alignment tool.

As predicted by the Signal P 4.1 software, these proteins have an N-terminal signal peptide marked in green. Protein domain searches did not reveal any known conserved domain.

